# Erythrocyte invasion-neutralising antibodies prevent *Plasmodium falciparum* RH5 from binding to basigin-containing membrane protein complexes

**DOI:** 10.1101/2022.09.23.509221

**Authors:** Abhishek Jamwal, Cristina E. Constantin, Sebastian Henrich, Wolfgang Bildl, Bernd Fakler, Simon J. Draper, Uwe Schulte, Matthew K. Higgins

## Abstract

Basigin is an essential host receptor for invasion of *Plasmodium falciparum* into human erythrocytes, interacting with parasite surface protein PfRH5. PfRH5 is a leading blood-stage malaria vaccine candidate and a target of growth-inhibitory antibodies. However, basigin is not alone on the erythrocyte surface. Instead, we show that it is exclusively found in one of two macromolecular complexes, bound predominantly to either plasma membrane Ca^2+^-ATPase 1/4, PMCA1/4, or monocarboxylate transporter 1, MCT1. PfRH5 binds to either of these complexes with a higher affinity than to isolated basigin ectodomain, making it likely that these are the physiological targets of PfRH5. PMCA-mediated Ca^2+^ export is not affected by PfRH5, ruling this out as the mechanism underlying changes in calcium flux at the interface between an erythrocyte and the invading parasite. However, our studies rationalise the function of the most effective growth inhibitory antibodies targeting PfRH5. While these antibodies do not reduce the binding of PfRH5 to monomeric basigin, they do reduce its binding to basigin-PMCA and basigin-MCT complexes. This indicates that the most effective PfRH5-targeting antibodies inhibit growth by sterically blocking the essential interaction of PfRH5 with basigin in its physiological context.

## Introduction

Malaria is still one of the most deadly parasitic diseases to affect humans, with *Plasmodium falciparum* as the causative agent of the most severe cases (Sato, 2021). The clinical symptoms of malaria occur as the merozoite developmental form of the parasite invades and replicates within human red blood cells (Venugopal et al., 2020). Vaccines or therapeutics which block erythrocyte invasion therefore have the potential to contribute to reduction and elimination of malaria.

Erythrocyte invasion is driven by a series of molecular interactions between merozoite ligands and their receptors on erythrocyte surfaces (Cowman et al., 2017). While many of these interactions are redundant, the binding of merozoite *P. falciparum* Reticulocyte Homologue 5 (PfRH5) to erythrocyte basigin is essential for invasion by all tested *Plasmodium falciparum* strains (Crosnier et al., 2011), making PfRH5 one of the most promising blood-stage malaria vaccine candidates. PfRH5 is part of the ternary PfRCR complex, containing PfRH5, PfCyRPA and PfRIPR, and each component is essential for invasion and a target of growth-inhibitory antibodies (Reddy et al., 2015; Wong et al., 2019).

Vaccination of human volunteers with PfRH5 elicits strain-transcending anti-malarial antibodies (Payne et al., 2017) and, in a human challenge model, reduces the rate of parasite growth (Minassian et al., 2021). Structural studies of PfRH5 have revealed that basigin binds towards one of the tips of the “kite-like” structure of PfRH5. Structures of PfRH5 in complex with Fab fragments of monoclonal antibodies have identified a number of important epitopes across PfRH5, including those for human and mouse growth-inhibitory antibodies (R5.004, R5.016, 9AD4 and QA1) (Alanine et al., 2019; Wright et al., 2014), as well as the epitope for an antibody which potentiates the effects of growth-inhibitory antibodies (R5.011) (Alanine et al., 2019). This insight has guided development of improved PfRH5-containing vaccines (Campeotto et al., 2017). However, mysteries remain, with the most effective growth-inhibitory antibodies, 9AD4 and R5.016, not able to block basigin binding. How do they prevent invasion?

Basigin is a type I transmembrane protein of the immunoglobulin (Ig) superfamily, with the most common isoform presenting an ectodomain consisting of two highly-glycosylated extracellular Ig domains (Muramatsu, 2016), both of which contact PfRH5 (Wright et al., 2014). More recently, the transmembrane segment of basigin has been shown to mediate formation of heteromeric complexes which contain basigin and either monocarboxylate transporters (MCTs) or plasma membrane calcium ATPases (PMCAs) (Muramatsu, 2016; Supper et al., 2016). MCTs are involved in proton-coupled exchange of lactate or pyruvate across plasma membranes (Felmlee et al., 2020). Both MCT1 and MCT4 interact with basigin in multiple eukaryotic cell lines (Kirk et al., 2000), affecting their function and stability (Kirk et al., 2000) and MCT1 is the primary subtype in human erythrocytes (Juel et al., 2003). PMCAs also play a role in membrane transport, with four subtypes of PMCA removing Ca^2+^ from the cell cytosol to maintain intracellular Ca^2+^ homeostasis and modulate cell signalling (Stafford et al., 2017). Either basigin, or its close paralog neuroplastin, are essential subunits of PMCA, influencing expression and function of PMCAs in rodent neurons, with human erythrocyte PMCAs complexed with basigin (Schmidt et al., 2017). Both basigin-MCT1 and neuroplastin-PMCA1 complexes have been structurally characterised, revealing interactions mediated by transmembrane helices (Gong et al., 2018; Wang et al., 2021), with the PfRH5 binding site on the basigin ectodomain remaining accessible.

Proteomics studies have confirmed that PMCA1, PMCA4 and MCT1 are all expressed on human RBC plasma membrane (Ravenhill et al., 2019) and, intriguingly, the actions of both MCTs and PMCAs have been linked to severe malaria (Bedu-Addo et al., 2013; Mariga et al., 2014; Timmann et al., 2012). In the case of MCT1, cerebral malaria is associated with increased lactate concentrations, which may damage the blood-brain barrier. However, it is hard to envisage a mechanism by which modulation of MCT1 function by PfRH5 during the rapid process of erythrocyte invasion could contribute to malaria symptoms. In contrast, PMCA polymorphisms have been linked to development of severe malaria in a genome-wide association study, suggesting that calcium homeostasis across the erythrocyte membrane may be linked to the outcomes of malaria. Indeed, spikes of increased intra-erythrocytic calcium have been observed during the parasite invasion process, and these are dependent on the interaction between PfRH5 and basigin (Volz et al., 2016; Weiss et al., 2015), raising the question of whether localised blockage of PMCA-mediated calcium efflux may be involved in these calcium spikes. In this study, we therefore asked whether PfRH5 can interact with basigin which is complexed with MCT1 and PMCAs, whether this modulates PMCA function and whether this explains the function of growth-inhibitory antibodies.

## Results

### Human erythrocyte basigin is primarily found in complex with PMCAs and MCTs

We first aimed to determine whether basigin is mostly free when extracted from human erythrocyte membranes, or is predominantly in complex with PMCAs or MCTs. We solubilised human erythrocyte ghosts in DDM/CHS, fractionated this extract by size-exclusion chromatography and used Western blotting to assess where basigin, PMCAs and MCT1 elute (Figure 1a). Probing individual fractions with a basigin-binding antibody revealed a broad band indicating the presence of heterogeneously glycosylated basigin. Fractions rich in basigin also showed strong signal for both PMCAs and MCT1, indicating that the majority of basigin co-migrates with PMCAs and MCT1 (Figure 1a). The three components predominantly eluted in a region of the migration profile between standards of mass 157kDa and 200kDa, compatible with the mass of detergent-solubilised basigin-MCT1 and basigin-PMCA complexes (Figure 1a). No basigin was found in later fractions, in the region of the profile which would correspond to the mass of a single basigin in a detergent micelle. These data suggest that basigin is predominantly found in complexes which co-elute with MCT1 and PMCA.

**Figure 1:**
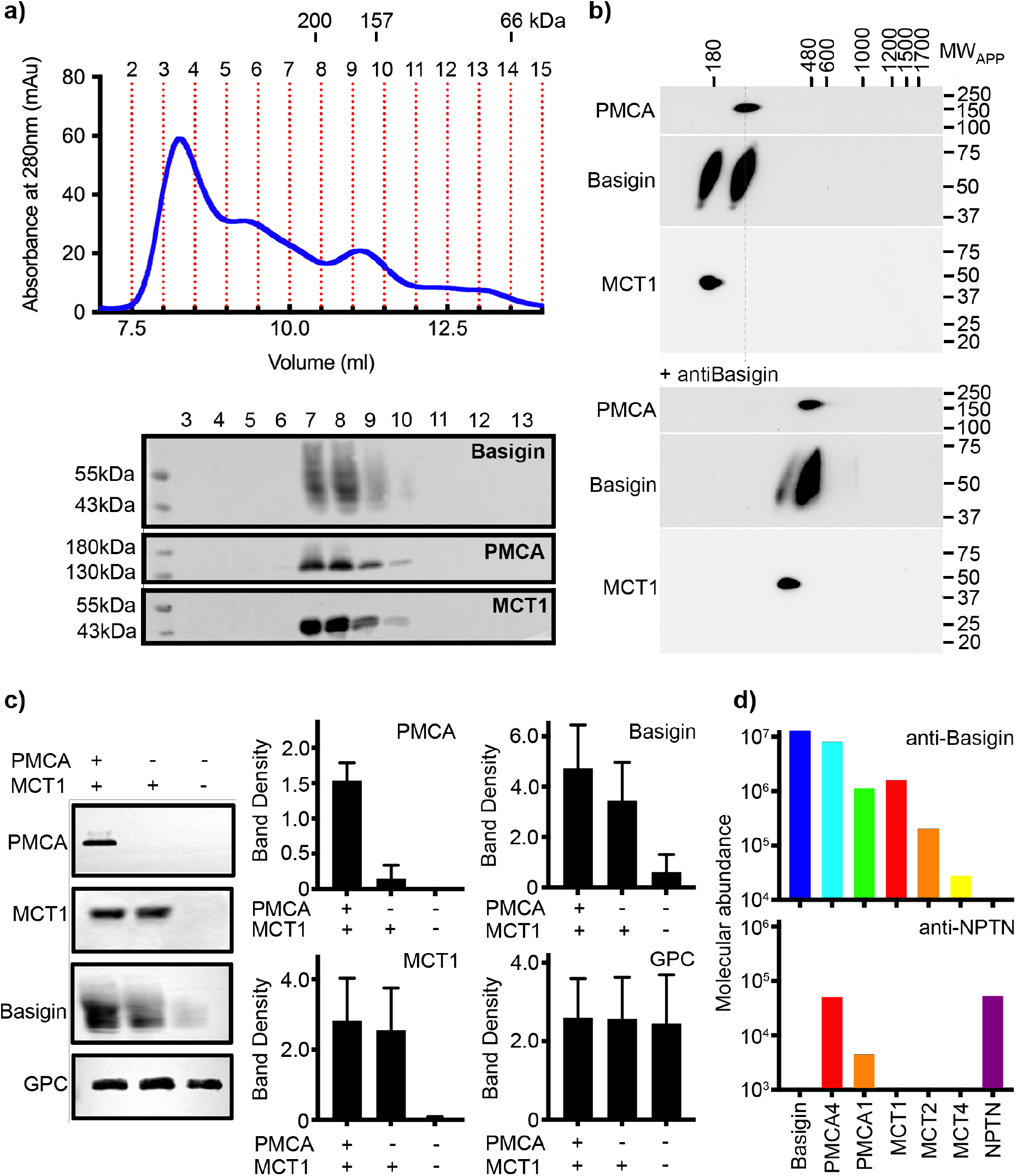
Basigin from human erythrocytes is found in complex with PMCA or MCT1. **a)** The upper panel shows the trace from size exclusion chromatography of DDM/CHS solubilized ghost membrane proteins fractionated on a Superdex 200 increase 10/300 column. The elution volumes of molecular weight standards are indicated above the trace. Fractions collected are demarcated by vertical red dotted lines. The lower panel shows Western blots of fractions 2 to 14. Representative blots for basigin (40-65 kDa, upper panel), PMCA (~ 138 kDa, middle panel) and MCT1 (~ 45 kDa, lower panel) show that all three proteins co-elute predominantly in fractions 7 to 9. **b)** Western blot analysis of 2D blue native PAGE/SDS-PAGE separations of human erythrocyte membrane solubilisates before (upper panel) and after pre-incubation with anti-basigin antibody (lower panel). Blot membranes were stained with antibodies specific for PMCA1/4, basigin and MCT1. Markers of apparent complex size indicate the positions of known mitochondrial respiratory chain (super)complexes (Schagger and Pfeiffer, 2000) run in a separate gel lane. Binding of the antibody led to a full size-shift of both PMCA1/4-basigin and MCT1-basigin complexes, whereas no signal of free basigin could be observed in the low molecular weight range, even after overexposure of the blot. **c)** The left-hand panel shows representative western blot images depicting sequential depletion of PMCA (upper panel) and MCT1 (upper-middle panel). Depletion of both transporters leads to reduced basigin levels, while levels of glycophorin C are unaffected in each fraction, again confirming that basigin is in complex with PMCA or MCT1. The remaining panels show densitometry plots obtained from inverted images of the western blots. Mean integrated band densities are shown with error bars as the standard error of the mean (n=3). **d)** Bar diagram depicting molecular abundances (abundance_norm_spec values) of the indicated proteins in depleting affinity purifications with anti-basigin and anti-neuroplastin (NPTN) antibodies from mildly solubilized human erythrocyte membranes as determined by mass spectrometry. Note the high co-purification efficiency of basigin interactors with AT2B4 (PMCA4) being its major partner. The abundance for all proteins was 0 after affinity purification with an IgG control.

To more closely investigate the association of basigin with PMCAs and MCT1, we performed antibody shift assays in two different settings. First, we analyzed antibody shifts of basigin-MCT1 complexes by size exclusion chromatography (Supplementary Figure 1a). A basigin-rich solubilised ghost extract was applied to a size exclusion column either in the presence or absence of an MCT1-targeting monoclonal antibody and the resultant fractions were probed on Western blots for MCT1, basigin or a glycophorin C control. While there was no shift of glycophorin C, the addition of the MCT1-specific antibody caused a shift in elution profile of the majority of the MCT1 and ~53% of basigin, suggesting them to be in complex (Supplementary Figure 1a). However, ~47% of the basigin did not shift. This identifies pools of basigin which are found in complex with and without MCT1. Next, we performed antibody shift experiments on detergent-solubilised human erythrocyte membrane extracts resolved by 2D blue native PAGE (Figure 1b). Western blotting these 2D gels and staining with basigin, PMCA and MCT1 antibodies, indicated two separate populations of basigin-containing complexes, one of which co-migrated with PMCA and one of which comigrated with MCT1. Inclusion of an antibody which quantitatively binds to basigin shifted both of these populations to a higher molecular weight. Signals at the appropriate molecular weight for free basigin were not observed in either the presence or absence of the basigin-binding antibody (Figure 1b).

In parallel, we conducted a depletion experiment. PMCAs and MCT1 were sequentially depleted from solubilized ghosts. PMCAs were first depleted by affinity capture on a calmodulin resin, while a monoclonal antibody was used to subsequently immunodeplete MCT1. Western blotting was used to assess the fraction of PMCA, MCT1 and basigin depleted at each stage and these were quantified by densitometry (Figure 1c). Sequential depletions of both PMCA and MCT1 resulted in almost complete elimination of basigin from solubilized ghost membrane proteome, showing that ~78% of basigin is complexed with one of these transporters. Densitometric measurements also showed greater reduction in basigin levels upon MCT1 depletion, with ~55 % of basigin complexed with MCT1 and ~27 % of basigin complexed with PMCA. Comparison with the outcomes of 2D blue native PAGE suggests that free basigin observed here most likely occurs as a result of disruption of protein complexes during immuno-depletion experiments.

As there are multiple PMCA subtypes in human cells, we next used mass-spectrometry analysis to identify which subtypes co-purify with basigin from human erythrocytes. This revealed PMCA4 to be the major partner of basigin in human erythrocytes, while also identifying PMCA1 (Figure 1d). Also identified were MCTs, with MCT1 as the primary subtype and MCT2 and 4 also detected (Figure 1d).

Collectively, these data show that native basigin in human erythrocytes is found only in heteromeric complexes, either with PMCA (predominantly PMCA4) or MCT1. As the interaction between RH5 and basigin has been shown to be essential for erythrocyte invasion by *Plasmodium falciparum*, this raises the question of whether RH5 can bind to either the basigin-PMCA and/or basigin-MCT1 complexes.

### PfRH5 binds to the basigin-PMCA complex with greater affinity than for monomeric basigin

We next investigated the binding of PfRH5 to basigin-PMCA complexes. The structure of the neuroplastin-PMCA1 complex shows the neuroplastin ectodomain to be presented above the extracellular loops of PMCA, with the surface of neuroplastin equivalent to that used by basigin to binding to PfRH5 exposed, making it likely that PfRH5 can bind to PMCA-bound basigin (Figure 2a) (Gong et al., 2018). To test this, we purified basigin-PMCA complexes from detergent solubilized human erythrocyte ghost cells using calmodulin affinity chromatography. Elution fractions showed a prominent band at ~ 138 kDa on a Coomassie stained SDS-PAGE gel, corresponding to the mass of PMCA (Supplementary Figure 2a). Western blotting confirmed the co-purification of basigin with PMCA (Supplementary Figure 2a). Basigin and PMCA eluted as a single peak on a size exclusion column, with an elution profile matching a 200kDa standard, suggesting complex formation (Figure 2b). The purified protein complex showed Ca^2+^-dependent ATP hydrolysis activity, confirming that active basigin-PMCA complexes had been isolated (Supplementary Figure 2b).

**Figure 2:**
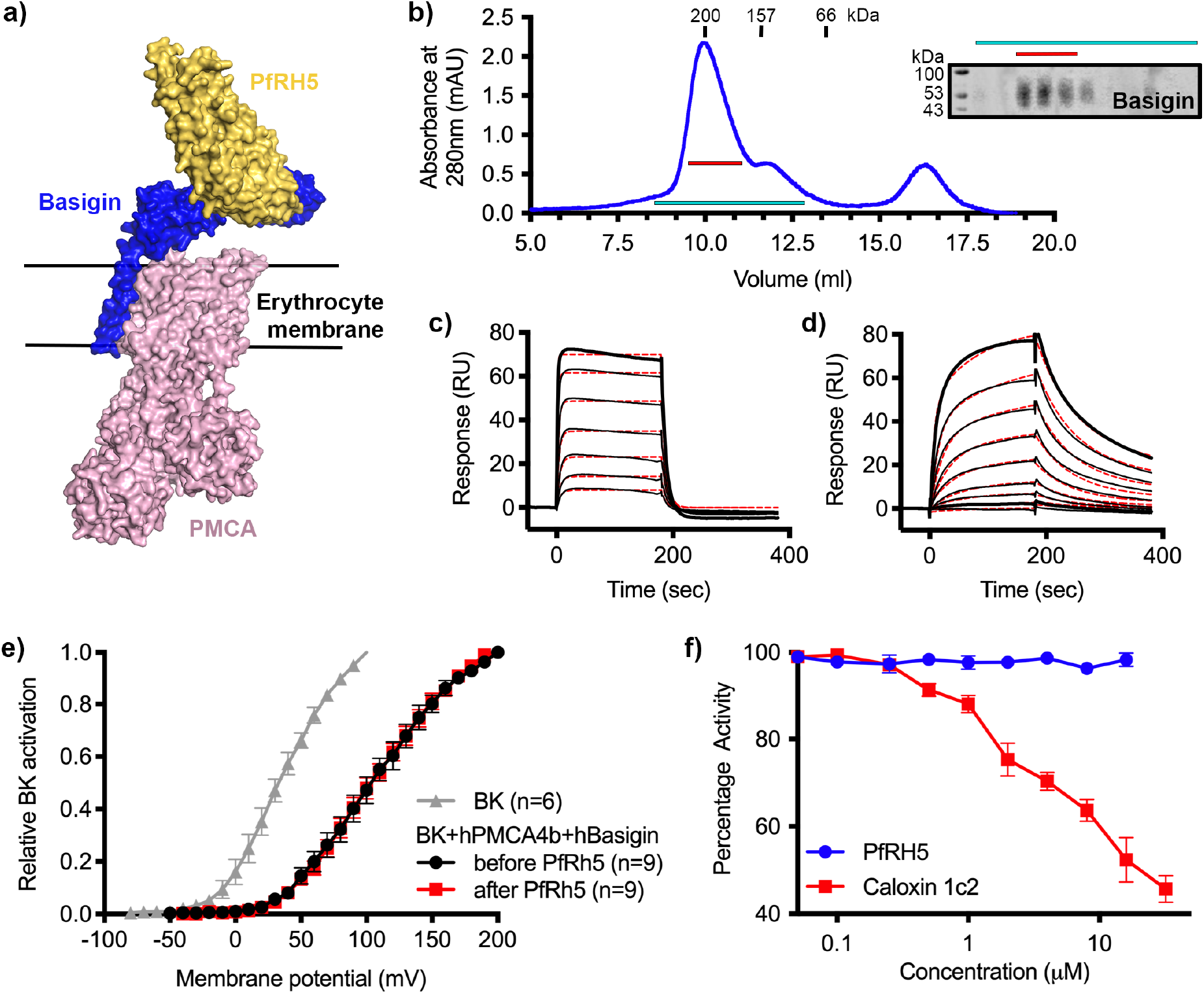
PfRH5 binds to basigin-PMCA without affecting calcium pumping activity. **a)** A structural model showing a complex of basigin (blue) and PMCA (pink) (based on PDB: 6A69 (Gong et al., 2018)) onto which the complex of PfRH5 (yellow) and basigin (blue) (PDB:4U0Q (Wright et al., 2014)) has been docked. **b)** A size exclusion profile obtained for calmodulin affinity chromatography-purified PMCA from human erythrocyte ghosts separated on a Superdex 200 increase 10/300 column. The inset shows a Western blot probed with an anti-basigin monoclonal antibody indicating co-migration of basigin with PMCA. **c)** An SPR sensogram showing the binding of a concentration series of basigin ectodomain (twofold dilutions from 4 μM) to immobilised PfRH5. Black lines show data and dashed red lines show fitting to a 1:1 binding model. **d)** An SPR sensogram showing the binding of a concentration series of basigin-PMCA (two-fold dilutions from 400 nM) to immobilised PfRH5. Black lines show data and dashed red lines show fitting to a two-state binding model. The derived rate and affinity constants are presented in Supplementary Tables 1 and 2. **e)** Activation curves of BK_Ca_ channels recorded in CHO cells expressing BK_Ca_ channels alone (grey), or cells also transfected with human PMCA4b and basigin with (red) and without (black) addition of PfRH5 at 2 μM concentration. Error bars show standard error of mean. **f)** Concentration-response curves for the inhibition of Ca^2+^-ATPase activity of purified basigin-PMCA complex by known inhibitor caloxin 1c2 (red) and PfRh5 (blue). Each data point represents the mean and error bars represent the standard deviation (n=3).

Binding of purified basigin-PMCA complexes to PfRH5 was measured by surface plasmon resonance (SPR) analysis. PfRH5 was immobilized on a streptavidin-coated chip and either 0.4 μM basigin-PMCA or 4 μM of basigin ectodomain were injected. Basigin ectodomain bound, generating a sensogram of a similar shape to that reported previously (Supplementary Figure 3a, Figure 2c) (Crosnier et al., 2011; Wright et al., 2014). Basigin-PMCA complexes bound PfRH5 with a sensogram profile indicating a slower off-rate (Supplementary Figure 3a, Figure 2d). To confirm that these responses are specific, we preincubated surface immobilised PfRH5 with monoclonal antibody R5.004, which occludes the basigin binding site (Alanine et al., 2019). This blocked the binding of both basigin ectodomain and basigin-PMCA to PfRH5, as expected (Supplementary Figure 3a).

To measure binding kinetics, we produced PfRH5 carrying an N-terminal biotin acceptor peptide tag. This was captured at low density on a streptavidin-coated SPR chip and binding was studied by injecting 2-fold dilution series of either basigin-PMCA or basigin ectodomain (Figure 2c and d, Supplementary Figure 3b and c). Kinetic binding data for basigin ectodomain could be fitted to a 1:1 binding model with an affinity of 0.78 μM (χ^2^ = 2.44), in good agreement with previous studies (Figure 2c) (Crosnier et al., 2011; Wright et al., 2014). In contrast, the kinetic data for the basigin:PMCA complexes did not fit well to a 1:1 binding model (χ^2^ = 7.71) (Supplementary Figure 3b), but the fit improved significantly (χ^2^ = 1.85) when using a two-state binding model (Figure 2d, Supplementary Figure 3b, Supplementary Tables 1 and 2). This model is consisted with an interaction which involves two separate, successive events, such as an initial lower affinity capture event, followed by a second binding event which increases the overall affinity, or a conformational change which follows an initial binding event. PfRH5 therefore bound to basigin-PMCA ~9-fold more tightly than to the isolated basigin ectodomain, primarily due to a lower dissociation rate constant (Supplementary Tables 1 and 2).

### PfRH5 binding does not modulate calcium transport mediated by basigin-PMCA complexes

We next investigated whether PfRH5 binding can modulate the activity of PMCA, perhaps accounting for changes in calcium flux at the junction between the erythrocyte and invading merozoite. To test this, we used an established system in which current recordings from BK_Ca_-type Ca^2+^-activated K^+^ channels were used as a readout of apparent intracellular Ca^2+^ concentration (Schmidt et al., 2017). The expression of PMCA4b and basigin, together with BK_Ca_, in this system caused reduced intracellular Ca^2+^ concentrations, as demonstrated by a right shift in the BK_Ca_ activation curve. We next added PfRH5 at a concentration of 2 μM and this caused no further change in the BK_Ca_ activation curve, indicating that PfRH5 binding did not modulate PMCA-basigin function and did not change the intracellular Ca^2+^ concentration (Figure 2e). As PfRH5 is normally part of the three component PfRCR complex, we also added PfRCR at a concentration of 1 μM to the same system. Here too, we observed no effect on intracellular Ca^2+^ concentration (Supplementary Figure 2c). Finally, we studied the effect of PfRh5 on Ca^2+^-dependent ATPase activity of purified basigin-PMCA using colorimetry. While the extracellular peptide inhibitor caloxin 1c2 inhibited calcium-dependent ATP hydrolysis at high micromolar concentrations, the addition of PfRh5 did not alter the ATPase activity of PMCA (Figure 2f). We therefore see no evidence that PfRH5 binding affects the activity of the basigin-PMCA calcium-pump.

### PfRH5 binds to the basigin-MCT1 complex with greater affinity than for monomeric basigin

As up to 55 % of erythrocyte basigin is associated with MCT1, we also tested whether the basigin-MCT1 complex can bind to PfRH5, as the PfRH5-binding site of basigin is exposed in the structure of the basigin-MCT1 complex (Wang et al., 2021) (Figure 3a). We first studied the binding of PfRH5 to solubilized ghosts and their depleted forms, using surface plasmon resonance. Flowing solubilized ghost membrane extract over a chip surface coated with PfRH5 produced a binding response (Figure 3b). This was reduced ~22% after depletion of PMCA-containing complexes, and the binding response was nearly absent for samples depleted for both PMCAs and MCT1 (Figure 3b). This suggests that basigin in complex with either MCT1 or PMCAs can bind to PfRh5.

**Figure 3:**
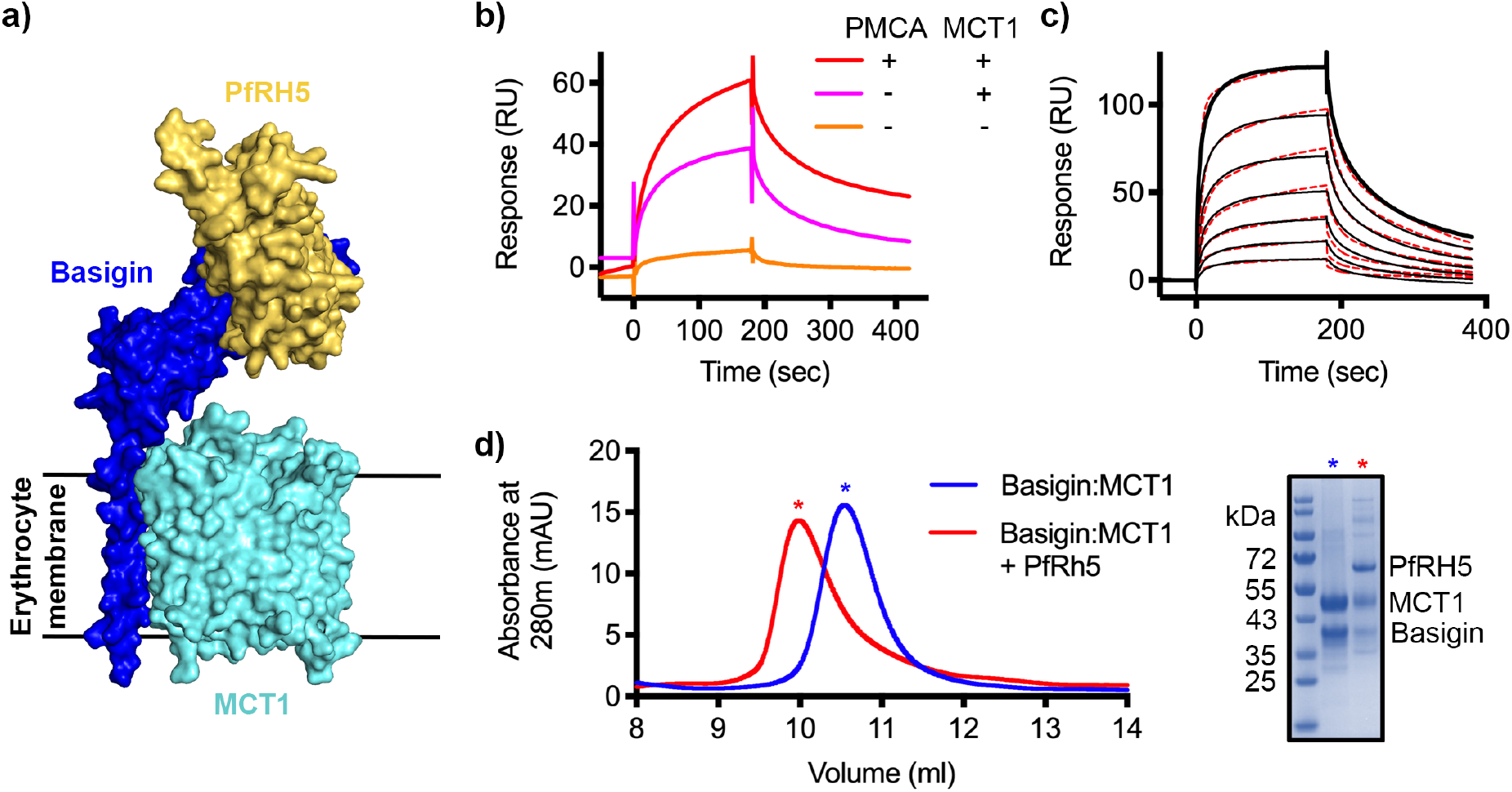
PfRH5 binds to basigin-MCT1. **a)** A structural model showing a complex of basigin (blue) and MCT1 (cyan) (based on PDB: 6LYY (Wang et al., 2021)) onto which the complex of PfRH5 (yellow) and basigin (blue) (PDB:4U0Q (Wright et al., 2014)) has been docked. **b)** SPR traces after flowing detergent solubilised erythrocyte membrane basigin-rich fractions (red) and membrane fractions depleted for PMCA (pink) and both PMCA and MCT1 (orange) over a PfRH5-coated surface. **c)** An SPR sensogram showing the binding of a concentration series of basigin-MCT1 (two-fold dilutions from 1600 nM) to immobilised PfRH5. Black lines show data and dotted red lines show fitting to a two-state binding model. **d)** Purified PfRh5 and basigin-MCT1 were assayed for complex formation through size exclusion chromatography using a Superdex 200 increase 10/300 column. The elution profile of basigin-MCT1 alone (blue) and in the presence of PfRh5 (red) are shown. The inset SDS-PAGE gel shows the protein species present in the fractions indicated by stars in the elution trace.

We next expressed the basigin-MCT1 complex in Sf9 cells and tested its binding to PfRH5. To measure the affinity of the interaction, mono-biotinylated PfRH5 was coupled to an SPR chip before injection of a 2-fold dilution series of purified basigin-MCT1 complex. Again, the kinetic data could be best described by a two-state binding model (χ^2^ = 3.09) with a K_D_ value of 93 nM (Figure 3c, Supplementary Figure 3c, Supplementary Tables 1 and 2). One of the predictions of a two-state binding model is that longer periods of incubation of PfRH5 with the basigin-MCT1 complex allow a larger fraction of interacting species to form stable complexes due to completion of the second binding event, and lead to slower off rates. When we injected basigin-MCT1 over a chip decorated with biotinylated PfRH5, we indeed observed slower off-rates to be associated with longer incubation times, confirming a two-state interaction (Supplementary Figure 3d). We also mixed basigin-MCT1 with PfRH5 for analysis by size exclusion chromatography, and again observed formation of a stable complex (Figure 3d). Therefore, PfRH5 binds to both basigin-PMCA and basigin-MCT1 with similar affinities, both more tightly than the interaction with the isolated basigin ectodomain.

### Growth-inhibitory antibodies sterically block binding of PfRH5 to basigin-PMCA and basigin-MCT1 complexes

While antibodies which occlude the basigin binding site of PfRH5 are growth-inhibitory, some of the most potent growth-inhibitory antibodies do not prevent PfRH5 from binding to basigin, raising questions about their mode of action. We therefore tested the ability of a panel of structurally characterised antibodies to prevent PfRH5 from binding to basigin ectodomain, basigin-PMCA and basigin-MCT1. The panel consisted of one growth-inhibitory antibody, R5.004, which prevents basigin binding, and two potent growth-inhibitory antibodies, R5.016 and 9AD4, which do not prevent basigin binding. Also tested were non-inhibitory antibodies, R5.011 and R5.015 which do not block basigin binding.

In accordance with previously published findings, only R5.004 prevented the binding of basigin ectodomain to PfRH5 immobilised on an SPR chip (Figure 4a, Supplementary Figure 4a). In contrast, both R5.016 and 9AD4 also reduced binding of the basigin-PMCA complex to immobilised PfRH5 by >90% (Figure 4b, Supplementary Figure 4b). Indeed R5.016 inhibited binding of basigin-PMCA to PfRH5 in a concentration dependent manner, with concentrations of 100 nM or above producing nearly 90 % inhibition (Supplementary Figure 4d). Both R5.016 and 9AD4 also reduced the binding of the basigin-MCT1 complex to immobilised PfRH5, with 100 nM of either antibody reducing binding by more than 50 % (Figure 4c, Supplementary Figure 4d). In contrast, neither of the non-inhibitory antibodies tested reduced the binding of basigin-PMCA or basigin-MCT1 to PfRH5.

**Figure 4:**
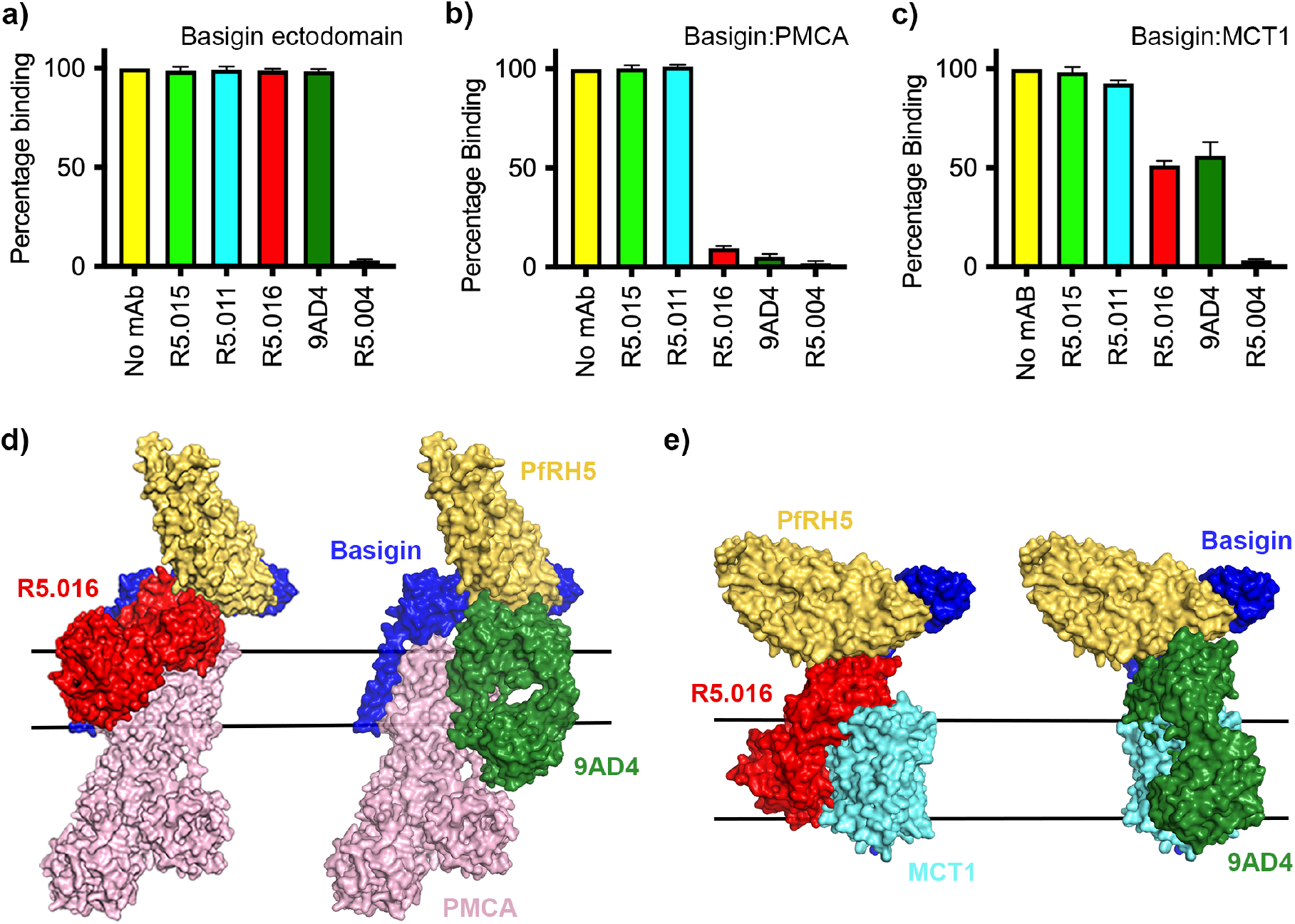
Neutralising monoclonal antibodies targeting PfRH5 prevent its binding to basigin-PMCA and basigin-MCT1 complexes. Five different PfRH5-binding monoclonal antibodies were tested for inhibition of the binding of PfRH5 to **a)** basigin ectodomain, **b)** basigin-PMCA complex and **c)** basigin-MCT1 complex. Data show is the mean (n=3) and error bars represent standard error of mean. **d)** Structural models showing a complex of basigin (blue) and PMCA (pink) (based on PDB: 6A69 (Gong et al., 2018)) onto which the complex of PfRH5 (yellow) and basigin (blue) (PDB:4U0Q (Wright et al., 2014)) has been docked. Onto this model has been docked either the complex of PfRH5 (yellow) bound to the Fab fragment of R5.016 (red, PDB:6RCV (Alanine et al., 2019)) or of 9AD4 (green; 4U0R (Wright et al., 2014)). **e)** Structural models showing a complex of basigin (blue) and MCT1 (cyan) (based on PDB: 6LYY [(Wang et al., 2021)) onto which the complex of PfRH5 (yellow) and basigin (blue) (PDB:4U0Q (Wright et al., 2014)) has been docked. Onto this model has been docked either the complex of PfRH5 (yellow) bound to the Fab fragment of R5.016 (red, PDB:6RCV (Alanine et al., 2019)) or of 9AD4 (green; 4U0R (Wright et al., 2014)).

To rationalise the mechanism of inhibition, we docked crystal structures of PfRh5 complexed with basigin (Wright et al., 2014), R5.016 (Alanine et al., 2019) and 9AD4 (Wright et al., 2014) Fab fragments onto the neuroplastin-PMCA (Gong et al., 2018) structure (Figure 4d) and the basigin-MCT1 (Wang et al., 2021) structure (Figure 4e) to generate composite models. In both cases, binding of R5.016 and 9AD4 to PfRH5 would physically occlude its binding to basigin in the basigin-PMCA or basigin-MCT1 complexes, either through direct clashes with MCT1 or PMCA, or through a clash with the plasma membrane. These data therefore support a model in which neutralising antibodies prevent PfRH5 from binding to basigin, in the context of macromolecular basigin-containing complexes.

## Discussion

Numerous studies have now confirmed that the interaction between PfRH5 and basigin is required for the invasion of *Plasmodium falciparum* into human erythrocytes (Crosnier et al., 2011). To date, no strain of *Plasmodium falciparum* has been identified which can take an alternative, basigin-independent, invasion route. Blocking this interaction also has proven therapeutic potential to prevent malaria, with antibodies against both PfRH5 and basigin being able to inhibit invasion (Alanine et al., 2019; Douglas et al., 2014; Wright et al., 2014; Zenonos et al., 2015), with vaccination (Douglas et al., 2015) or passive transfer of neutralising antibodies (Douglas et al., 2019) leading to protection of aotus monkeys from parasite challenge and with human vaccination slowing the onset of parasitaemia (Minassian et al., 2021). Nevertheless, mysteries remain. First, the functional consequences of the PfRH5-basigin interaction are still unclear. What are the downstream outcomes of the interaction and why are they essential for invasion? Second, some of the most potent human and mouse PfRH5-targeting monoclonal antibodies which block parasite invasion do not directly block the PfRH5-basigin interaction. How do they function?

To answer these questions, we started by investigating the immediate molecular context of basigin in human erythrocytes. Various studies in different cell types have shown that basigin forms heteromeric complexes with either PMCAs or MCTs (Muramatsu, 2016). We therefore used a combination of depletion, purification, 2D blue native PAGE and mobility shift experiments to show that the majority of basigin in human erythrocyte membranes is found in complex with either PMCA or MCT1 (Figure 1). While ~20% of basigin was not depleted with either PMCA or MCT1 (Figure 1c), all basigin migrated at a high molecular weight on a size exclusion column and we did not observe free basigin in 2D blue native PAGE experiments, suggesting that in techniques which more carefully preserve complex formation, basigin is bound to PMCAs or MCTs (Figure 1b). The detection of free basigin in depletion experiments may then be due to disruption of protein complexes during depletion experiments, or due to basigin in complex with subtypes or splice isoforms of PMCA or other subtypes of MCTs not efficiently depleted. Thus, the vast majority of basigin in human erythrocytes is found within larger protein complexes, with consequences for its recognition by PfRH5 and its role in *Plasmodium* invasion.

Intriguingly, both MCT1- and PMCA-bound basigin bind more tightly to PfRh5 than to isolated basigin ectodomain. Indeed, fitting of kinetic binding data from SPR showed that, while the binding of PfRH5 to basigin ectodomain fits a simple 1:1 binding mode, its binding to basigin-PMCA and basigin-MCT1 shows biphasic kinetics, which fit most closely to a two-state binding model. This is further supported by data which shows that PfRH5 binds to basigin-MCT1 such that increasing association times correlate with decreasing dissociation rates, suggesting an initial lower affinity interaction, which ‘matures’ into a higher affinity interaction over time. While kinetic fitting data cannot be used to strictly derive a molecular mechanism, this is compatible with an initial lower affinity interaction between PfRH5 and basigin, followed by a subsequent interaction which stabilises the complex, perhaps between PfRH5 and another part of the basigin-PMCA or basigin-MCT1 complexes.

Our demonstration that PfRH5 can bind to basigin-PMCA and basigin-MCT1 next led us to investigate whether PfRH5 binding can modulate the functions of these two transporters. MCT1 is a lactate transporter, and we were not able to envisage a mechanism by which altered lactate transport could play an essential role in the context of the ~20 second-long process of erythrocyte invasion. However, PfRH5 is known to be essential for a spike of increased calcium concentration at the merozoite-erythrocyte junction (Volz et al., 2016; Weiss et al., 2015) and basigin binding has been shown to alter PMCA function (Schmidt et al., 2017). Could modulation of basigin-PMCA function block calcium efflux and cause this spike? Our data do not support this hypothesis, with no change in basigin-PMCA function observed, either in the presence of PfRH5 or of the PfRH5-PfCyPRA-PfRIPR complex. How PfRH5 modulates the erythrocyte to facilitate invasion is therefore still unclear. However, the demonstration that erythrocyte basigin is primarily found in large multimeric complexes will provide clues for future research to unveil this mechanism.

Finally, our results reveal the mechanisms by which the most potent PfRH5-targeting neutralising antibodies can block invasion. Immunity to blood stage malaria is antibody-mediated and, while antibodies which target other aspects of the malaria life cycle use Fc-dependent mechanisms of immunity (Kurtovic et al., 2020; Teo et al., 2016), those which prevent invasion appear to function directly through blocking essential interactions (Alanine et al., 2019; Douglas et al., 2019; Wright et al., 2014). As a result, the outcome of *in vitro* growth-inhibition assays correlates well with protection due to PfRH5-targeting antibodies (Douglas et al., 2019). It was therefore a mystery why the most effective PfRH5-targeting neutralising antibodies from both humans (R5.016) (Alanine et al., 2019) and mice (9AD4) (Douglas et al., 2014; Wright et al., 2014) do not interfere with the PfRH5-basigin interaction. Studying basigin in the context of its membrane protein binding partners provides an answer, with both R5.016 and 9AD4 inhibiting the binding of PfRH5 to basigin-PMCA and basigin-MCT1 complexes. While this inhibition is strong in the context of a detergent micelle, we would expect it to be even stronger in the membrane context, with both R5.016 and 9AD4 modelled to angle directly towards the membrane plane. Our studies therefore highlight the importance of understanding the molecular context of basigin in understanding PfRH5 function and its inhibition, and open new directions to understand why the PfRH5-basigin interaction is so crucial for erythrocyte invasion.

## Materials and Methods

### Ghost preparation from human erythrocytes

400-450 ml of packed human red blood cells (RBCs) were purchased from a regional hospital. RBCs were lysed in 10 volumes of hypotonic buffer (10 mM Tris-Cl, pH 8.0 and 1mM EDTA and 1mM PMSF) at 4 ºC for 2 hours. The lysate was first centrifuged at 1500 g for 30 mins to remove cellular debris and then circulated through four omega T-series centramate 0.1 sq. ft tangential filter flow unit (100 kDa, PALL) to generate 300-350 ml of concentrated retentate of RBC membranes. The retentate was spun at 100,000 g for 45 mins at 4 °C and the bulk haemoglobin was very carefully pipetted from the top without disturbing the pellet. To remove residual haemoglobin, retrieved membrane pellets were resuspended in hypotonic buffer and centrifuged at 100,000 g for 20 mins at 4 °C. This process was repeated at least four times or until white to light pink ghosts were obtained. Ghosts corresponding to 100 ml of packed RBCs were stored in 50 ml of buffer containing 20 mM HEPES-Na pH 7.2, 150 mM NaCl, 10 % glycerol supplemented with complete protease inhibitor cocktail EDTA-free (Roche) at −80 °C until further use.

#### Fractionation and analysis of solubilized ghosts by size exclusion chromatography

0.5 ml of ghost membranes were solubilized by adding 10 % DDM-CHS (10:1 w/w ratio) solution to final concentration of 1.2 % and incubated for 1 hour at 4 °C with gentle stirring. Solubilized membranes were spun at 100,000 g for 45 mins at 4 °C, and the supernatant was injected into a Superdex 200 increase 10/300 GL (cytiva) column calibrated with known molecular weight standards (GE healthcare). Size exclusion chromatography was then performed at a flow rate of 0.5 ml/min in a buffer containing 20 mM HEPES pH 7.2, 150 mM NaCl and 0.02% DDM/CHS where proteins eluting from the column were collected as discrete fractions of 500 μl each. 10-15 μg protein from each fraction was separated on a 10% SDS-PAGE gel and blotted on a PVDF membrane using Bio-Rad turbo transfer pack and apparatus. Membranes were blocked with 6 % TBS-milk subsequently probed with 1:500 mouse antihuman PMCA IgG (5F3, Santa Cruz Biotechnology), 1:1000 mouse anti-human CD147 IgG (Biolegend) and 1:1000 mouse anti-human MCT1 IgG (Santa Cruz Biotechnology) overnight at 4 °C. Washes were conducted in TBS and TBS-T-20 0.05 % and the membrane was then incubated with anti-mouse IgG 1:20000 for 45 mins at room temperature. After final washes, blots were developed and visualized using ECL reagent SuperSignal^TM^ West Atto (Thermo Scientific) and i-Bright imager (Life Technologies).

### Mobility shift using an antibody against MCT1

200 μl of 0.1-0.15 mg of fractions of DDM-CHS solubilized human ghost proteins rich in basigin were incubated with 20 μg of mouse anti-human MCT1 IgG and incubated for 30 mins on ice. Following the incubation, the sample was injected into a Superdex 200 increase 10/300 GL column and eluted at 0.5 ml/min as fractions of 500 μl. 10-15 μg protein from each SEC fraction from samples with or without MCT1 antibody were separated by 12% SDS-PAGE, prior to Western blotting. Detection of MCT1 and basigin was performed on separate blots, using 1:1000 of polyclonal rabbit anti-human MCT1 IgG (Cell Signalling) and 1:1000 mouse anti-human CD147 IgG (Biolegend) followed by HRP-conjugated secondary IgG antibody of specific host type. Proteins were visualized with ECL reagent SuperSignal^TM^ West Atto using i-Bright imager (Life Technologies).

### Depletion of native PMCA and MCT1

0.20 mg of solubilized ghost proteins were incubated with 30 μl of calmodulin (CaM) resin for 45 mins at 4 ºC to deplete native PMCA. Beads were removed by centrifugation at 250 g for 5 mins. 50 μg of supernatant was set aside for analysis and the remaining sample was mixed with 10 μg of mouse anti-human MCT1 IgG and incubated for 30 mins on ice. Following the incubation, 50 μl of protein G resin was added to the mixture and was further incubated for 2 hours on a rotating platform in a cold room. Supernatant from this depleted mixture was collected by centrifugation to remove beads. 10-15 μg of protein from solubilized ghost and each depleted fraction was separated on 12 % SDS-PAGE for side-by-side analysis using Western blotting. Blots were developed and visualized as described above.

### Affinity purification from human erythrocytes for mass spectrometry analysis

Membranes were prepared from human erythrocytes by hypotonic lysis (1:50 (v:v) in 1 mM EDTA/EGTA, 10 mM Tris-HCl pH 7.4 + protease inhibitors) followed by ultrasound treatment (2x 20 pulses, duty cycle 50, output 1 (Branson Sonifier 250)) and ultracentrifugation for 10 min at 150,000 x g. Membrane pellets were resuspended at 11 mg/ml (as determined by Bradford assay, Bio-Rad) in 20 mM Tris-HCl pH 7.4. 2 mg of membrane were solubilized in 2 ml ComplexioLyte 47 ready-to-use detergent buffer (Logopharm) supplemented with 1 mM EDTA/EGTA and protease inhibitors. After ultracentrifugation for 10 min at 250,000 x g, the solubilisate was precleared by incubation with 15 μg bead-immobilized control IgG (Millipore 12-370) for 1 h at 4°C. Two 0.5 ml aliquots of the supernatant were then mixed with 5 μg of bead-immobilized anti-basigin (Origene TA501189) and anti-neuroplastin (R&D Systems AF7818) antibodies, respectively. After 2 h incubation at 4°C, beads were washed twice for 10 min in ComplexioLyte 47 washing buffer (Logopharm) and eluted with 2 x 10 μl of denaturing buffer (non-reducing Laemmli buffer + 8M urea, 2x 10 min at 37°C). The eluates were then supplemented with 100 mM DTT, run on an SDS-PAGE gel and silver-stained. Lanes were split into high- and low-MW range sections and subjected to standard in-gel tryptic digestion for subsequent LC-MS/MS analysis. Target depletion in these affinity purifications was verified by Western blot analysis (not shown).

### Mass spectrometry and protein quantification

LC-MS/MS analysis was carried out in positive ion mode on an LTQ Orbitrap XL mass spectrometer (Thermo Scientific, Germany; CID fragmentation of the five most abundant new precursors per scan cycle; singly charged ions rejected; dynamic exclusion duration 30 s) equipped with a split-based UltiMate 3000 HPLC (Dionex/Thermo Scientific, Germany; “short” gradient) as detailed in (Schmidt et al., 2017). Peak lists were extracted from MS/MS spectra using ‘msconvert.exe’ (ProteoWizard; https://proteowizard.sourceforge.io/; v3.0.6906). After an initial database search precursor mass (m/z) values were corrected by linear shifting of their median offset. Final searches against the UniProtKB/Swiss-Prot database (human, release 2022_02) using Mascot Server 2.6.2 (Matrix Science Ltd, UK) were restricted to +-5 ppm (peptide mass tolerance) and used the following parameters: Acetyl (Protein N-term), Carbamidomethyl (C), Gln->pyro-Glu (N-term Q), Glu->pyro-Glu (N-term E), Oxidation (M), Propionamide (C), Phospho (ST) and Phospho (Y) as variable modifications, fragment mass tolerance ±0.8 Da, one missed tryptic cleavage allowed.

Label-free quantification of proteins was carried out as described (Muller et al., 2016). In brief, fullscan MS data was processed with MaxQuant v1.6.3 (https://www.maxquant.org) to obtain mass calibrated peptide signal intensities (peak volumes, PVs). Their elution times in the evaluated datasets were pairwise aligned and then assigned to peptides based on their m/z and elution time (obtained either directly from MS/MS-based identification or indirectly from identifications in parallel datasets) using in-house developed software. Matching tolerances were ± 2 ppm and ± 1 min. Abundance_norm_spec values reflecting the molecular abundance of proteins were calculated as the sums of all assigned and protein isoform-specific PVs divided by the number of MS-accessible protein isoform-specific amino acids (Bildl et al., 2012).

### 2D antibody shift assay

Blue native (BN) gel-based antibody shift analysis was carried out as described in (Schmidt et al., 2017). 2 mg of human erythrocyte membrane were solubilized and centrifugated as described for mass spectrometry analysis. 600 μl of the solubilisate was mixed with anti-basigin antibodies (8.4 μg, TA501164 (9H5) + TA501189 (10E10), both Origene) for 2 h and 600 μl remained untreated. Samples were concentrated on a sucrose cushion (200 μl 20% sucrose in ComplexioLyte 47 + protease inhibitors and 300 μL 50% sucrose in 750 mM aminocaproic acid and 50 mM BisTris pH 7.0) by ultracentrifugation at 400,000 x g for 1.5 h. 300 μl of both sucrose phases, together with 120 μl of solubilized rat brain mitochondria (serving as size marker), were separated overnight on a 2-12% BN-PAGE gradient gel. The lanes were excised, equilibrated in 2x Laemmli buffer with 8 M urea and resolved on second dimension 10% SDS-PAGE gels that were blotted on PVDF membranes. The latter were cut, blocked with 3% skim milk powder in PBS / 0.05% Tween-20 and stained with anti-PMCA1/4 (Alomone Labs ACP-005), anti-basigin (Proteintech 11989-1AP) or anti-MCT1 (Santa Cruz Biotechnology SC-365501) followed by HRP-conjugated secondary IgG antibodies specific for respective host species (Santa Cruz Biotechnology) and ECL Prime captured by film. Total protein stains of these western blot membranes (SYPRO Ruby Protein Blot Stain, Bio-Rad) were used for proper alignment of blot signals and visualization of size marker complexes.

### Expression and purification of BAP-tagged PfRh5

A construct was generated consisting of PfRH5 with an N-terminal biotin acceptor peptide (BAP) and a C-terminal C-tag, inserted into the pExpreS2.1 vector (ExpreS2ion Biotechnologies, Hørsholm, Denmark). This was transfected into Schneider 2 (S2) cells and after 24 hours zeocin was added to a final concentration of 0.05 mg/ml. A polyclonal cell line was selected over three weeks and was then expanded to higher volumes for isolation and purification.

1.5 litres of cell supernatant were concentrated tenfold, and buffer exchanged to Trisbuffered saline (TBS) pH 7.8 using four omega T-series centramate 0.1 sq. ft tangential filter flow unit (3 kDa, PALL). The buffer-exchanged supernatant was loaded onto a CaptureSelect™ C-tagXL pre-packed column (Thermofisher Scientific). The column was washed with 50 ml of TBS, pH 7.8 and PfRh5 was eluted with buffer containing 20 mM Tris-Cl pH 7.8 and 2mM MgCl_2_ as six fractions of 500 μl each. Each fraction was quantified at 280 nm using a nano-spectrophotometer (Thermofisher Scientific) and fractions of concentration 0.5 mg/ml or more were subjected to separation on a Superdex 200 increase 10/300 GL column (cytiva) for further purification.

### Purification of basigin-PMCA complex from ghosts

100 ml of frozen human ghost cells were thawed in a water bath at 20 ºC. Ghosts were then solubilized by adding DDM/CHS solution to final concentration of 1.2 % at 4 ºC under gentle stirring for 1 hour. Non-solubilized material was removed by centrifugation at 100,000 g for 30 mins and CaCl_2_ was added to the supernatant to final concentration of 0.25 mM. 1ml of calmodulin resin (MERCK) pre-equilibrated in wash buffer containing 20 mM HEPES-Na pH 7.2, 300 mM NaCl, 10% glycerol, 0.25 mM CaCl_2_, and 1.2% DDM/CHS was added to the supernatant mixture and incubated at 4 ºC under gentle stirring. After 1 hour, the resin was transferred to a gravity flow column and washed with 50 ml of equilibration buffer containing 0.02% instead of 1.2% DDM/CHS. Finally, protein bound to calmodulin resin was eluted in 8-10 fractions of 250-300 μl each in total 2.4 ml of elution buffer containing 20 mM HEPES-Na pH 7.2, 150 mM NaCl, 1 mM EDTA and 0.02 % DDM/CHS. Identity and purity of eluted proteins was assessed by coomassie staining on 10 % SDS PAGE followed by Western blotting with mouse anti-human basigin IgG (Biolegend). Fractions were pooled and concentrated using Amicon centrifugal filter units 100 kDa (Merck) to separate on a Superdex 200 increase 10/300 GL column equilibrated (GE healthcare) with buffer containing 20 mM HEPES-Na pH 7.2, 150 mM NaCl and 0.02% DDM/CHS to check homogeneity or integrity of affinity purified complex. Using this procedure, a total of 20-35 μg of basigin-PMCA complex was obtained.

### Recombinant expression and purification of basigin-MCT1 complex

Synthetic genes (GeneArt) encoding full-length human MCT1 (uniprot ID: P53985) and human basigin (uniprot ID: P35613-2) were cloned individually into pFastBac vector (Invitrogen) for recombinant expression in Sf9 cells. MCT1 was fused to a C-terminal 6X-His tag, whereas basigin was not tagged. Bacmids were generated in DH10Bac cells (Life Technologies) and purified by isopropanol precipitation using a miniprep kit (Qiagen). First generation baculoviruses (P1) were produced by transfecting Sf9 cells with individual bacmids at a cell density of 1 x 10^6^ cells/ml and subsequently amplified to produce third generation viruses (P3). Expression of the complex was induced by adding an equal number of viral particles from both MCT1 and basigin containing P3 viruses to Sf9 cells (~2.5-3.0 x 10^6^ cells/ml). After 48 hours cells were harvested and resuspended in lysis buffer (25 mM Tris pH 8.0, 150 mM NaCl, 10 % glycerol and EDTA-free complete protease inhibitor) and lysed using a Dounce homogeniser sonication (60% amplitude, 3 secs ‘on’ and 9 secs ‘off’ pulse for 1:30 mins) on ice. Lysed homogenate was centrifuged at 3000 g for 20 mins to remove cell debris, and the obtained supernatant was spun at 100,000 g for 45 mins at 4 ºC to isolate Sf9 plasma membranes. Isolated membranes homogenised in lysis buffer were solubilized with 1.2% DDM/CHS at 4 °C for 1 hour. After centrifugation at 100,000 g for 45 min, the supernatant was incubated with Ni-NTA resin (Qiagen) at 4 °C for 1 hour. The resin was then passed through on a gravity flow column, which was washed with 25 mM Tris pH 8.0, 300 mM NaCl, 10% glycerol, 15 mM imidazole, and 0.02% DDM/CHS. The protein was eluted in lysis buffer supplemented with 400 mM imidazole. Eluted protein complex was further purified on a Superdex 200 increase 10/300 GL column, and a total of 0.2-0.3 mg could be obtained from 500 ml Sf9 cell culture.

### Surface plasmon resonance affinity measurements

SEC purified PfRH5 was biotinylated enzymatically using *E. coli* biotin ligase to transfer a single biotin residue. Experiments were performed at 25 °C on a Biacore T200 instrument (GE Healthcare) using a CAP chip (Biotin CAPture kit (cytiva)) in a buffer containing 20 mM HEPES-Na pH 7.5, 150 mM NaCl, 1mM EDTA, 0.02% DDM/CHS and 1mg/ml salmon sperm DNA. Purified analyte proteins were equilibrated in the SPR buffer using a PD-5 column prior to the experiment. The sensor surface was first coated with oligonucleotide coupled streptavidin following manufacturer’s instructions. 50-70 RUs of monobiotinylated PfRH5 were captured on the chip to quantify interactions with basigin-PMCA and basigin-MCT1 complexes, whereas 300-350 RUs were immobilized for basigin ectodomain binding. Binding measurements were performed at 30 μl/min by injecting two-fold concentration series from 400 nM to 1.56 nM for basigin-PMCA and 1.6 μM to 6.25 nM for basigin-MCT1 complexes. In contrast, a twofold series from 4 μM to 7.81 nM was used to generate a kinetic profile for the basigin ectodomain. Association was measured for 180 s followed by dissociation for 300 s and after each binding cycle, the sensor chip surface was regenerated by injecting 10μl of 6M guanidium-HCl and 1M NaOH pH 11.0 mixed in a 4:1 ratio. The data was processed using BIA evaluation software version 1.0 (BIAcore, GE healthcare) and response curves were double referenced by subtracting the signal from both reference cell and averaged blank injections. Kinetic constants were calculated using global analysis, fitting 1:1 Langmuir model (A + B = AB) for basigin-ectodomain, and a two-state model involving conformation change (A+B=AB*=AB) for basigin-PMCA and basigin-MCT1 interactions with PfRh5.

### Surface Plasmon Resonance analysis using ghost proteins

SEC purified PfRH5 in 1X PBS was chemically biotinylated using EZ-Link^TM^ Sulfo-NHS-Biotin (Thermofisher Scientific) following manufacturer’s instructions. Basigin enriched ghost protein and its PMCA and MCT1 depleted versions were first exchanged into SPR buffer on a PD-5 column. Each Ghost fraction was then flowed individually over CAP chip surface coated with 400-450 RUs of chemically biotinylated PfRh5. The association phase was measured for 180 s, while the dissociation phase was observed for 300 s and the recorded response curves were double referenced by subtracting response data from reference surface and buffer blank.

### Surface plasmon resonance analysis of monoclonal antibody blocking

450-500 RUs of chemically biotinylated PfRH5 with was immobilized on flow cell 2 of the CAP chip. A mAb blocking experiment was then performed by injecting single concentration (typically 50 nM for each mAb) for 240 secs followed immediately by an analyte injection (2 μM basigin ectodomain or 0.2-0.4 μM basigin-PMCA or basigin-MCT1 complex) for 180 s on flow cells 1 and 2. Also, ten-fold dilution series of 10 μM - 0.1nM of R5.016 was applied for 240 s to gauge concentration dependent inhibition of the binding of PfRH5 to basigin-PMCA by R5.016. All sensogram curves were double referenced by subtracting responses from reference surface and blank injections.

### Measurement of Ca^2+^-ATPase activity

0.075-0.1μg of basigin-PMCA was incubated in a buffer containing 20 mM HEPES pH 7.2, 150 mM NaCl, 1 mM MgCl_2_, 0.5 mM EGTA and 0.1 mM ATP and 0.02 % (w/w) DDM/CHS. The reaction started by addition of 0.55 mM CaCl_2_, incubated for up to 30 min at 30 ºC. The reaction was stopped by addition of 1mM EGTA following which the inorganic phosphate (P_i_) released due to ATP hydrolysis was detected by using P_i_ color lock gold kit^TM^ (Abcam) and by measuring absorbance at 625 nm using a Spectra Max (Molecular Devices) colorimeter. The Ca^2+^-ATPase activity in the reaction mixture was calculated by subtracting the amount of P_i_ liberated in the tubes without Ca^2+^ and expressed in micromolar at different times points. The concentration of P_i_ in the reaction mixes was determined from a linear standard curve generated from P_i_ standard solution provided in the kit.

To study inhibition of activity, 0.1 μg of basigin-PMCA complex in 100 μl reaction buffer was pre-incubated with 0-32 μM of PfRH5 or caloxin 1c2 peptide in a 96-well microplate. The ATPase activity was stimulated by addition of 0.55 mM CaCl_2_ at 30 ºC. After 10 mins, the reaction was stopped with 1mM EGTA and colour was developed using P_i_ color lock^TM^ gold kit followed by absorbance measurements at 625 nm The inhibition of PMCA activity was inferred by comparing amount of P_i_ liberated in the reaction mixes containing PfRH5 or caloxin 1c2 to the reaction mix without any inhibitor.

### Electrophysiology

Chinese hamster ovary (CHO) cells were transiently transfected with cDNA coding for the mouse BK_(Ca)_ channel alpha subunit (Uniprot ID Q08460), PMCA4b (P23634) and Basigin (P35613). The cells were incubated at 37°C and 5% CO2 and measured 2-4 days after transfection. Whole-cell patch clamp recordings were performed at room temperature using a HEKA EPC 10 amplifier. The currents were low pass filtered at 3-10 kHz and sampled at 20 KHz. Leak currents were subtracted using a P/4 leak subtraction protocol with holding potential of −90mV and voltage steps opposite in polarity to those in experimental protocol. Serial resistance was 30-70% compensated using the internal compensation circuitry. The standard extracellular solution contained 5.8mM KCl, 144mM NaCl, 0.9mM MgCl_2_, 1.3mM CaCl_2_, 0.7mM NaH_2_PO_4_, 5.6mM D-Glucose and 10mM HEPES pH 7.4. Recording pipettes pulled from quartz glass had resistance of 2.5 – 4.5 MΩ when filled with internal solution containing 139mM KCl, 3.5mM MgCl_2_, 2mM DiBrBAPTA 5mM HEPES pH 7.3; 2.5mM Na2ATP, 0.1mM Na3GTP pH 7.3. CaCl_2_ was added to the internal solution to obtain 5 μM free [Ca^2+^]. PfRh5 and PfRCR were applied at final concentration of 2 and 1 μM respectively, for 3 to 10 minutes in the bath before recording.

Steady state activation of BK_Ca_ channels was determined using test pulses ranging from −50 to +200 mV (in 10 mV increment), followed by a repolarization step to 0mV. The conductance – voltage relations have been determined from the tail current amplitudes measured 0.5ms after repolarization to the fixed membrane potential (0 mV) and normalized to the maximum. For fitting, a Boltzmann function was used: g/g_max_ = g_max_/(1+exp((V_h_-V_m_)/k)), where Vh is the voltage required for half maximal activation and k the slope factor.

All the chemicals except DiBrBAPTA (Alfa Aesar) were purchased from Sigma.

### Data processing software

Image J was used for densitometric analysis of western blots. SPR Curves, SEC elution traces and other graphs were reproduced using Prism 9 software and structural figures were prepared with Pymol (Schroedinger).

## Acknowledgements

This work was funded by a Wellcome Investigator Award (20797/Z/20/Z) to MKH and was supported by grants of the DFG (SFB 746, TP 16, Fa 332/9-1) to B.F. and (SFB 746, TP 20) to U.S. The authors would like to thank David Staunton for support with biophysics data collection.

## Author Contributions

A.J., U.S. and M.K.H. conceived and planned the study and wrote the manuscript. A.J. conducted protein production, interaction analysis and biophysics studies. C.E.C. conducted electrophysiology experiments. S.H. conducted affinity purifications and 2D gel experiments. W.B. conducted MS analysis. B.F. provided infrastructure, project management and supervision for MS analysis. All authors designed experiments, analysed data and read and commented on the manuscript.

## Supplementary Figures

**Supplementary Figure 1:**
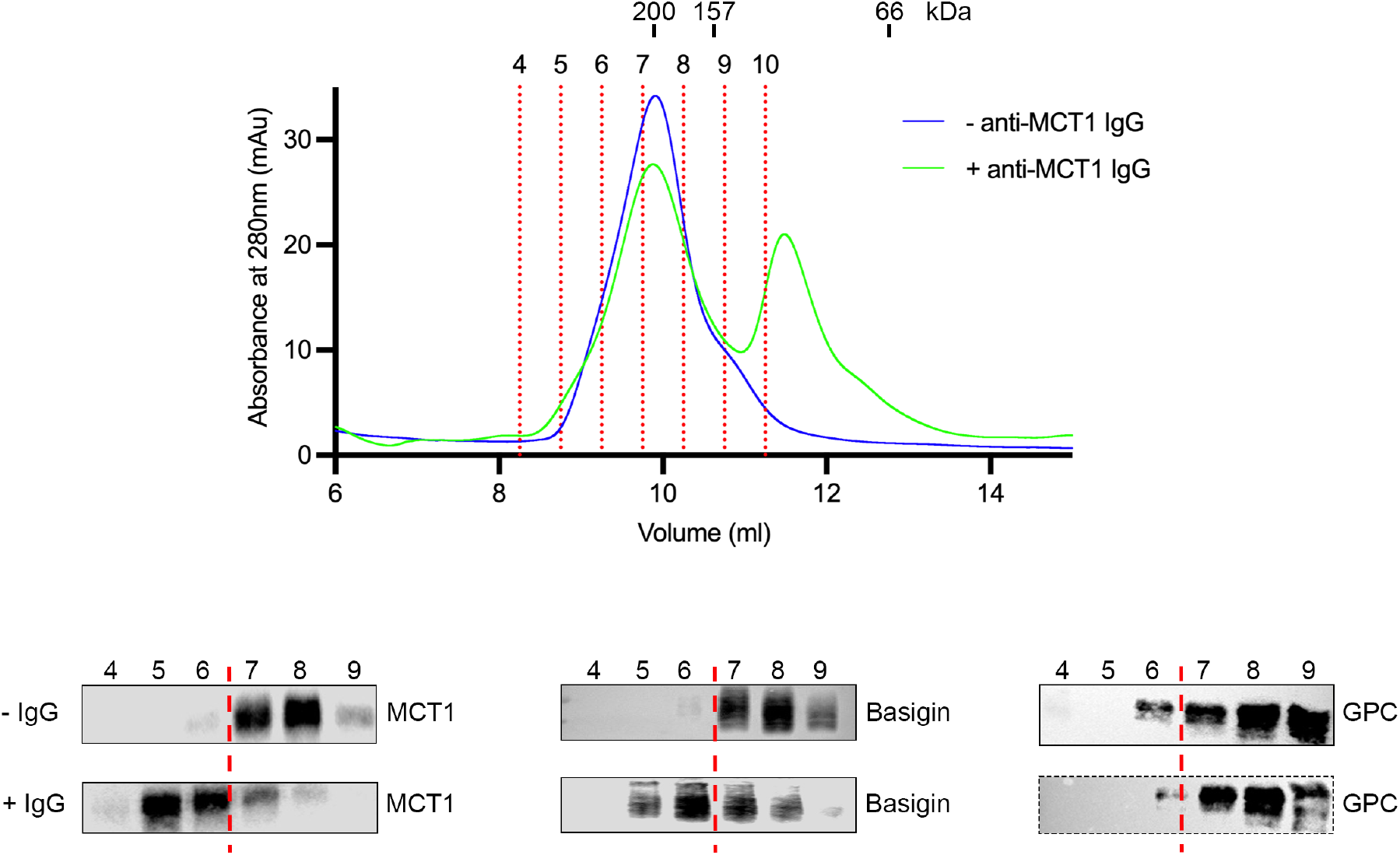
Mobility shift of basigin using an MCT1-targeting antibody. The top panel shows the outcome of separation of the basigin-rich fraction from DDM:CHS-solubilized ghost membranes on a Superdex 200 increase 10/300 column (blue). This was repeated in the presence of MCT1 antibody (green). Individual fractions are labelled and demarcated using red dashed lines. Above the trace are shown the elution profile of molecular weight standards. The lower panel shows Western blots probed with MCT1 (left), basigin (central) and GPC (right) for the fractions from the size exclusion profile, both without (top) and with (bottom) the MCT1 antibody. This shows mobility shift for both MCT1 and basigin, but not the glycophorin C (GPC) control.

**Supplementary Figure 2:**
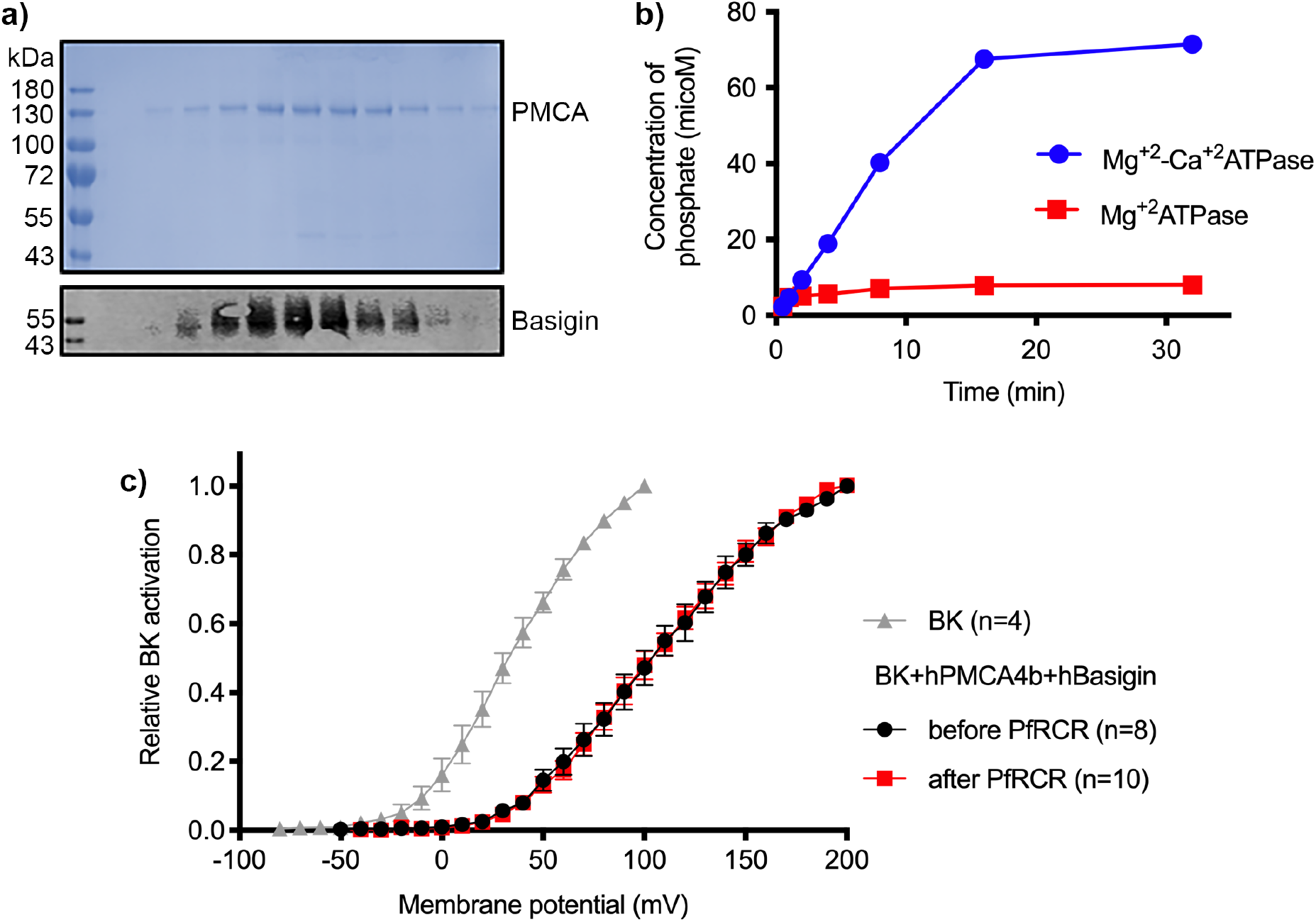
Purification and functional characterisation of basigin-PMCA. **a)** The upper panel shows Coomassie-stained elution fractions from the outcome of purification of PMCA from DDM:CHS solubilized erythrocyte ghosts using a calmodulin-affinity column. Sharp protein bands for PMCA can be observed running at ~130 kDa. The lower panel shows a Western blot of the same fractions probed with basigin monoclonal antibody. Basigin and PMCA co-elute. **b)** Measurement of phosphate release from ATP catalysed by basigin-PMCA in the absence (red) and presence (red) of Ca^2+^. **c)** Activation curves of BK_Ca_ channels recorded in CHO cells (grey), or cells also transfected with human PMCA4b and basigin with (red) and without (black) addition of the RCR complex (PfRH5, PfCyRPA and PfRIPR) at 1 μM concentration. Error bars show standard error of mean.

**Supplementary Figure 3:**
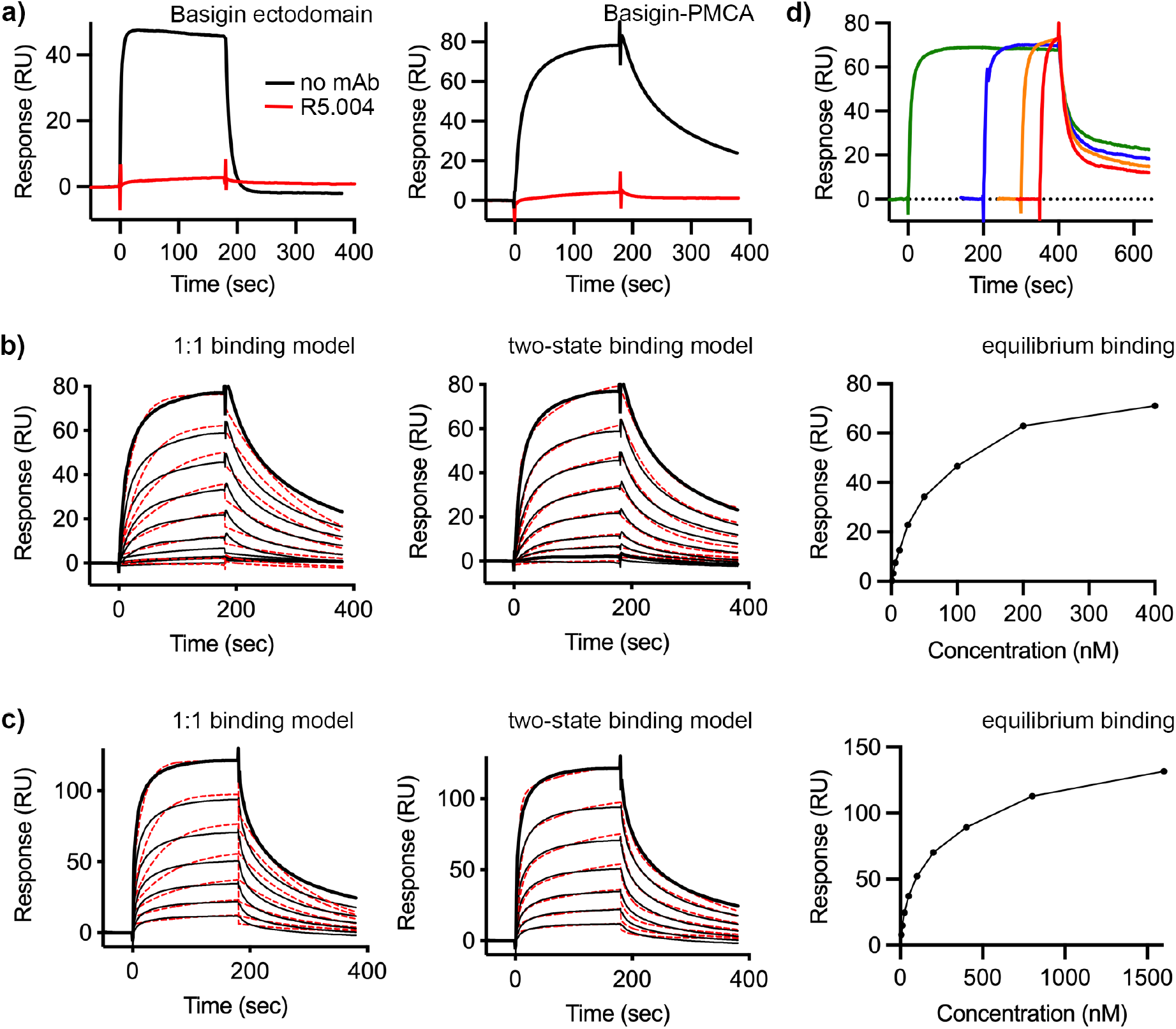
surface plasmon resonance analysis of binding of PfRH5 to basigin, basigin-PMCA and basigin-MCT1. **a)** The binding of basigin and basigin-PMCA to immobilised PfRH5 (at a concentration of 50 nM) in the presence (red) and absence (black) of monoclonal antibody R5.016. **b)** The binding of basigin-PMCA to immobilised PfRH5 (with a two-fold dilution series starting at 500 nM). Data were filled to a 1:1 binding model (left), a two-state binding model (centre) and equilibrium model (right, 92 nM affinity). **c)** The binding of basigin-MCT1 to immobilised PfRH5 (with a two-fold dilution series starting at 1600 nM). Data were filled to a 1:1 binding model (left), a two-state binding model (centre) and equilibrium model (right, 178 nM affinity). **d)** Analysis of the binding of basigin-MCT1 (at a concentration of 250 nM) to immobilised PfRH5. The four curves were measured with different association times, showing that increases association time leads to slow dissociation rates. In each case, data is shown as a black line and fitting curves are dashed red lines.

**Supplementary Figure 4:**
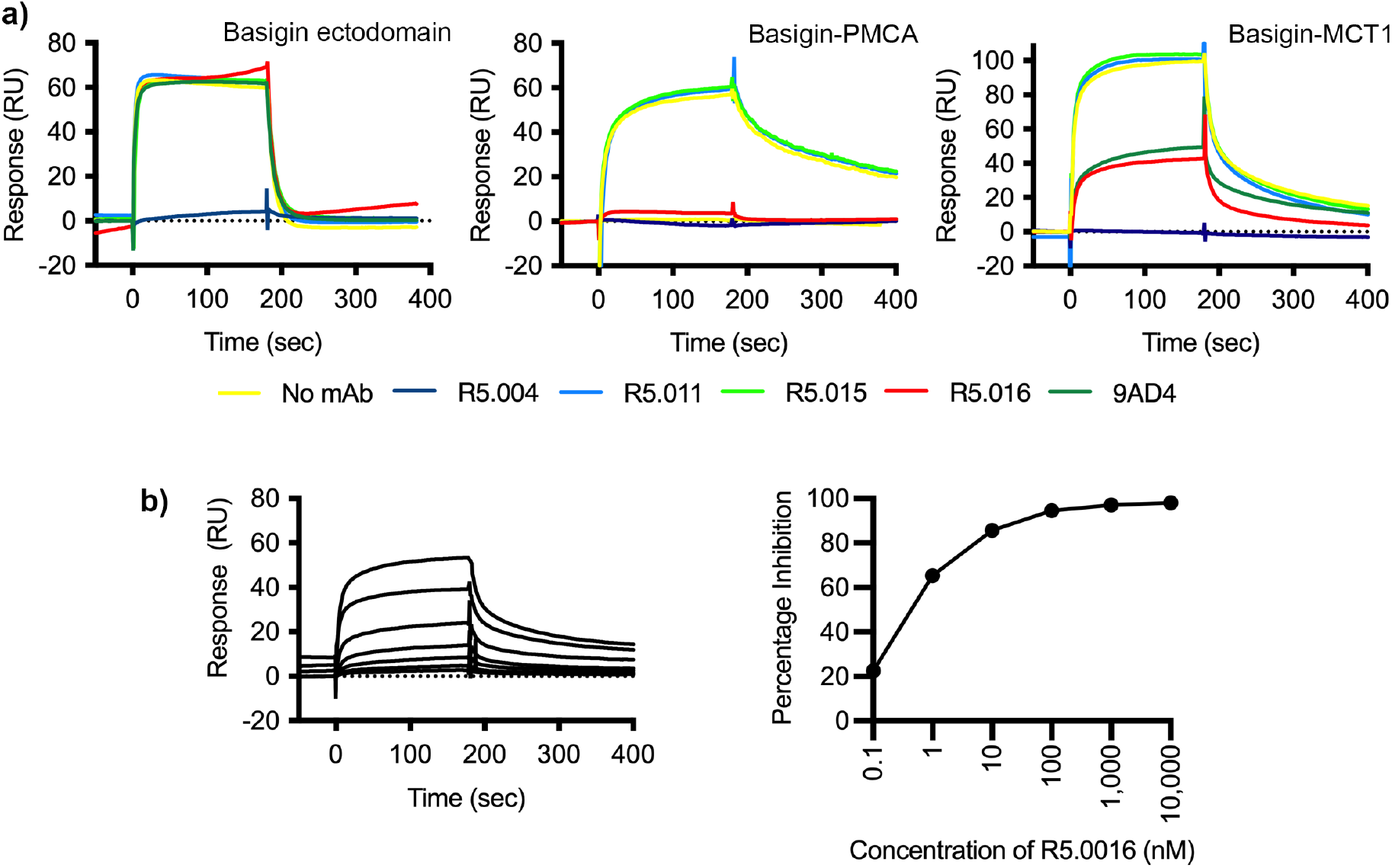
Neutralising monoclonal antibodies targeting PfRH5 prevent its binding to basigin-PMCA and basigin-MCT1 complexes. **a)** The impact of five different PfRH5-binding monoclonal antibodies was tested for inhibition of the binding of PfRH5 to basigin ectodomain (left), basigin-PMCA complex (centre) and basigin-MCT1 complex (right) by surface plasmon resonance analysis. **b)** Analysis of the dose response of inhibition of binding of basigin-PMCA to immobilised PfRH5 with different concentrations of R5.016. The left-hand panel shows the surface plasmon resonance traces while the right-hand panel shows the response plotted against R5.016 concentration.

**Supplementary Table 1.** Binding constants derived from surface plasmon resonance analysis by fitting sensograms with Langmuir 1:1 model

| Analyte | $k_a$ ($M^{-1}s^{-1}$) | $k_d$ ($s^{-1}$) | $K_D$ (M) | $R_{max}$ (RU) | $\chi^2$ (RU<sup>2</sup>) |
| --- | --- | --- | --- | --- | --- |
| Basigin ectodomain | $2.11 \times 10^5$ | 0.166 | $7.86 \times 10^{-7}$ | 79.98 | 2.44 |
| Basigin-PMCA | $9.55 \times 10^4$ | 0.007188 | $7.938 \times 10^{-8}$ | 85.01 | 7.71 |
| Basigin-MCT1 | $1.88 \times 10^5$ | 0.008743 | $7.357 \times 10^{-8}$ | 100.2 | 14.8 |

**Supplementary Table 2.** Binding constants derived from surface plasmon resonance analysis by fitting sensograms with two-state interaction model

| Analyte | $k_{a1}$ ($M^{-1}s^{-1}$) | $k_{d1}$ ($s^{-1}$) | $k_{a2}$ ($M^{-1}s^{-1}$) | $k_{d2}$ ($s^{-1}$) | $K_D$ (M) | $R_{max}$ (RU) | $\chi^2$ (RU<sup>2</sup>) |
| --- | --- | --- | --- | --- | --- | --- | --- |
| Basigin-PMCA | $3.48 \times 10^5$ | 0.0609 | $7.58 \times 10^3$ | $6.74 \times 10^{-3}$ | $8.78 \times 10^{-8}$ | 98 | 1.85 |
| Basigin-MCT1 | $4.26 \times 10^5$ | 0.0866 | $4.58 \times 10^3$ | $3.88 \times 10^{-3}$ | $9.31 \times 10^{-8}$ | 131 | 3.09 |

